# Investigation of Electrical Stimulation Effect on Intracellular Calcium Dynamics Using Locally Uniform Mesh Refinement

**DOI:** 10.1101/267294

**Authors:** Mohammad Omid Oftadeh

## Abstract

Modeling-based methods are conventionally exploited to simulate and predict the dynamics of a system in various fields of engineering. However, due to the intricacy of biological systems, these methods were rarely utilized in bioengineering until recently. By developing advanced computers with the ability to cope with enormous calculations and extending our knowledge about biological systems, modeling-based approaches have been adopted to manipulate biological systems. While utilizing the models to investigate the behavior of a system has some advantages including lower cost and time consumption, such methods were used for tissue engineering scarcely. Exploiting modeling-based methods to tissue engineering field requires developing and utilizing efficient computational methods to model gene regulation and signaling networks, which determine cell response to environmental changes and cellular fate.

In the present work, a novel spatio-temporal method was proposed, predicated on locally uniform mesh refinement. A benchmark comparison was performed with the previously used method and the results indicated the better performance of the incipiently adopted method. Besides, the model was utilized to investigate the effect of a popular differentiating stimulator, namely electrical stimulation, on mesenchymal stem cells as a type of stem cells widely used in tissue engineering. The results of the simulations demonstrated the puissance of such computational models in analyzing and predicting the effect of biochemical and biophysical perturbations on the cells and hence, their potential utility in tissue engineering.

## 1 Introduction

Regenerating defected or damaged tissues is one of the main challenges in the medical field, as current methods suffer from several shortcomings such as lack of tissue supply and immunogenicity (in the case of xenografting and allografting) and low remodeling (in the case of synthetic grafts) [1]. Tissue engineering is an emerging method concerning with the treatment of tissue deficiencies by stimulating cells (conventionally stem cells) to proliferate and differentiate and consequently, repair damaged site. To stimulate cell proliferation and differentiation, the effects of many biochemical and biophysical factors on the cells have been investigated [2]. Considering that several cell types such as neurons and heart muscle cells communicate with each other through electrical pulses, electrical stimulation (ES) is among these factors and it has been adopted for tissue engineering purposes in recent years, especially with the advent of conductive biocompatible scaffolds [3]. In addition to neurons and cardiac cells, ES has been applied for bone tissue engineering due to the piezoelectric effect of bone [4].

Generally, ES causes diverse effects on cells ranging from galvanotaxis (immigration toward anode or cathode), change in proliferation (whether increase or decrease), enhancement in growth factors expression (particularly vascular endothelial growth factor (VEGF), transforming growth factor (TGF)-β, epidermal growth factor (EGF), fibroblast growth factor (FGF)) and amending differentiation toward neurons, cardiac cells and osteoblasts by alteration in expression of different genes [3]. Despite the different and sometimes contradictory effects reported for ES impact on cells, virtually unanimously, all of them mediated through calcium ions [2, 3, 5–7] and distinctions between them mainly stem from the ES delivery setup [3]. In more details, applying ES causes the plasma membrane to depolarize and subsequently, activates voltage-gated calcium channels (VGCCs), which in turn, establishes Ca^2+^ influx to the cells [5]. Finally, upsurge in intracellular Ca^2+^ triggers downstream signaling pathways and activates components such as Ca^2+^/calmodulin-dependent protein kinase II (CamKII) [2]. In this vein, L-type calcium channels (LCCs) play the crucial role, as blocking them diminishes the ES effect on cells entirely [2, 5].

Ca^2+^ entrance into the cells via VGCCs additionally triggers more Ca^2+^ release from endoplasmic reticulum (ER) via inositol 1,4,5-trisphosphate receptor (IP_3_R) channels in the ER’s membrane. Ca^2+^ release from IP_3_R channels might have three different patterns based on their spatio-temporal behavior: the smallest events, called blips, are spatially restricted elevations in Ca^2+^ lasting less than a second and usually occur when a single channel opens which is widely assumed to be a stochastic event [8, 9]. The intermediate-size events, called Ca^2+^ puffs or spikes, occur when several spatially connected channels (called clusters) open simultaneously, customarily in response to an extracellular stimulus [10]. These events are local and last for a few hundred millisecond [8, 9]. The largest events, called waves, are global and include coordinated whole-cell Ca^2+^ oscillations caused by the opening of many channels in the ER membrane in response to external stimulus initiating large and long-lasting oscillations that sometimes diffuse to adjacent cells [8, 9]. The regularity, frequency and amplitude of Ca^2+^ puffs and waves determine the cell’s response to a stimulus in general [9] and ES in particular [11].

In this article a novel and efficient numerical method was proposed to model spatio-temporal dynamics of signaling networks. The method belongs to a class of numerical methods called spatial domain decomposition which, despite their efficiency have not been previously used in spatio-temporal modeling of signaling networks. After providing a benchmark comparison with the previous algorithms, the method was utilized to model the effect of ES on mesenchymal stem cells (MSCs) as a popular type of stem cells in tissue engineering. The model is only aimed to study the effect of ES on local Ca^2+^ pattern (Ca^2+^ puffs) and to evade nonessential intricacy, only the components whose crucial roles in mediating the effect of ES on the cells experimentally attested were incorporated into the model. The model is aimed to clarify the differences between the effect of various ES regimens used to drive MSCs to different cell lineages.

## 2 Methods

### 2.1 Electrophysiological Model of MSCs

To investigate the effect of ES on Ca^2+^ influx via plasma membrane, ion channels in the membrane of MSCs were modeled based on the Hodgkin-Huxley model. According to experimental investigations, human MSCs are not homogeneous and based on their excitation pattern divided into at least three types. Five different ionic channels have been identified in human MSCs: delayed rectifier-like hEAG1 channel, calcium activated potassium channel, tetrodotoxin (TTX)-sensitive sodium channel, L-type calcium channel and transient outward channel and each type of MSCs expresses a different combination of these channels [12, 13]. Since the main focus is on Ca^2+^ dynamics in this article, only one type of human MSC that expresses L-type calcium channels was investigated. In addition to L-type calcium current, other ionic currents, including tetrodotoxin (TTX)-sensitive sodium current and delayed rectifier-like hEAG1 and calcium activated potassium currents were also identified in this type of MSC (referred to as type C) [14]. The membrane potential of a single cell can be generally modeled as:

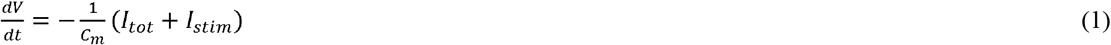

where V, t and C_m_ are voltage, time and membrane capacitance, respectively. I_stim_ is external stimulating current and I_tot_ is the total transmembrane current. In other words, I_tot_ is the sum of ionic currents passing through the membrane:

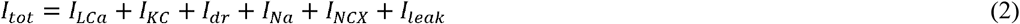

where I_Ca_, I_KC_, I_dr_, I_Na_, I_NCX_ and I_leak_ stand for L-type calcium, calcium activated potassium, delayed rectifier-like hEAG1, tetrodotoxin (TTX)-sensitive sodium, sodium-calcium exchanger and leakage currents, respectively. The required parameters to calculate equations (1) and (2) were estimated using whole-cell patch clamp results obtained in a previous work [12] (cf. supplementary information).

### 2.2 Stochastic gating of L-type Ca^2+^ channel (LCC)

Various modes of opening -closing have been identified for single LCC. The rates of conversion between various modes are dependent on several factors including the membrane potential and the presence of certain pharmaceuticals [15, 16]. The common mode of LCC gating is distinguishable with brief openings and longer closings and its open and close durations can be modeled as stochastic events with exponential distribution (with mean duration of opening and closing are τ=1.19 ms and τ=8.51 ms, respectively [16]). Here, another mode of gating characterized by long-lasting openings and brief closings was also considered in addition to normal mode. Similarly, opening and closing events for this mode of gating is consistent with exponential distribution (means are τ=19.26 ms and τ=0.36 ms, respectively [15, 16]). The conversion between this mode and normal mode is voltage-dependent and the equilibrium between two modes is inclined toward the mode with long-lasting openings by depolarization of membrane beyond 0 mV [15]. A simplistic kinetic scheme for the conversion can be proposed by considering gating modes as distinctive states that can be converted to each other by first order reactions [15]. Defining kl and kb rate constants respectively, for conversion of brief-opening to long-lasting-opening mode and vice versa results in the following equations:

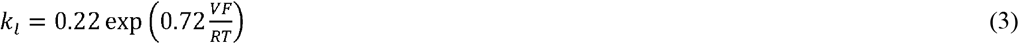

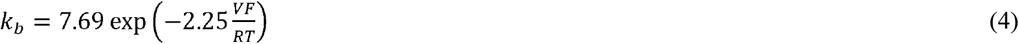

In the above equations, V, F, R and T are the membrane potential, Faraday constant, gas constant and temperature, respectively. Considering the conversion is at equilibrium, the equilibrium constant K_eq_=k_l_/k_b_ indicates the ratio of the presence of long-lasting-opening mode to normal mode and hence, the probability of triggering each mode can be calculated.

### 2.3 Stochastic gating model for IP_3_R channels

A stochastic version of DeYoung-Keizer model for IP_3_R channels was used. According to modified DeYoung-Keizer model, each channel has four subunits and each subunit has three binding sites: one binding site for activating Ca^2+^, one binding site for inhibiting Ca^2+^ and one binding site for IP_3_. An active channel has at least three activated subunits and each subunit is active when the binding site for IP_3_ and activating Ca^2+^ is occupied, while binding site for inhibiting Ca^2+^ is unoccupied [9]. Considering each binding site can be bound or unbound, there are eight different states for each subunit. However, to be more consistent with experimental results, an additional state was considered according to previous works [17]. Based on this model, an activated subunit becomes open only upon its transition to the additional state. All transitions between different states were modeled by Gillespie’s method for stochastic modeling of chemical reactions supposing that each cluster has nine channels.

### 2.4 Reaction-diffusion equations

To model spatio-temporal Ca^2+^ dynamics, a cubic intracellular subspace with 2 μm sides (approximate distance between ER membrane and the plasma membrane in MSC [18, 19]) was considered in which the upper and lower surfaces represent plasma membrane and ER membrane, respectively. Ca^2+^ entered into the subspace via LCC(s) and IP_3_R cluster and left the space through the pumps on the ER and plasma membranes. To mimic intracellular condition, both mobile and stationary Ca^2+^ buffers were considered. More details about reaction-diffusion formulation are proposed in the supplementary information.

### 2.5 Numerical methods

#### 2.5.1 Semi-discretization in space

Two methods for spatial discretization were used. The first method is based on non-uniform mesh refinement (NUMR) and previously proposed [20]. The second method was utilized for the first time to model intracellular signaling network and belongs to locally uniform mesh refinement (LUMR) class. The two methods were compared in terms of computation time and memory space.

For NUMR grid, finite element method (FEM) was used to discretize the reaction-diffusion equations. In general, the system of reaction-diffusion equations has the following form:

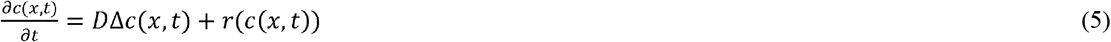

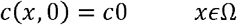

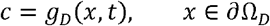

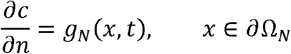

where c is the concentration field, D is the diffusion coefficient and r(c) represents the reaction term. Considering H^1^ as the space of square integrable functions, it is equipped with the following scalar product and corresponding norm:

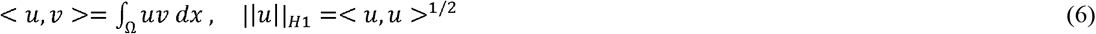

Multiplying equation (5) by 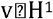 and integrating over Ω yield variational form of the system of equations:

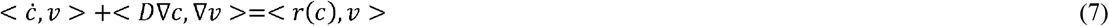

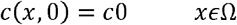

Replacing *v* with basis functions, equation (5) can be written in the following matrix form:

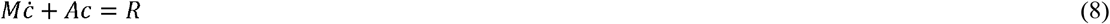

Where M and A are mass and stiffness matrices, respectively, and R is the reaction vector for each chemical specie.

To obtain the system of equations (8), the cubic space was discretized into non-uniform tetrahedral elements using a MATLAB mesh generator called DistMesh (a mesh generator that determines the location of nodes by solving equilibrium in a truss structure and then, connects them by Delaunay triangulation) [21]. To provide spatial adaptivity, the algorithm previously proposed [20] was used. The mass and stiffness matrices were calculated for the mesh and the reaction vector was estimated by numerical integration (quadrature). While no flux through lateral boundaries was considered, Ca^2+^ flowed into and out of the space through upper and lower boundaries (Neumann boundary) (cf. supplementary information).

The static-regridding method was used to construct LUMR grid. There are various static-regridding methods which are mainly based on common principles. The method used here is similar to the algorithm proposed by Trompert and Verwer [22] with some modifications.

First, the physical domain was discretized into a coarse grid using cubic cells. Integration over time-step Δt and over the coarsest (base) grid was performed. Δt might be equal to the base grid time-step, which was determined in a way that the solution became stable and sufficiently accurate. Afterward, the solution at each node was assessed to find approximate error at each cell. The edge of the cells with intolerable error bisected to obtain the refined mesh. In this vein, the distance between nodes in the refined mesh is half of the nearest coarse grid (parent grid). Then, integration was repeated over Δt and over the refined mesh taking smaller steps than the parent grid time-step. Integrating over the refined grid, two issues arise: how to obtain an initial value for the nodes in the refined mesh and how to treat boundaries. Regarding the first issue, if a node was present in the previous time-step, its last solution could be used as its initial value for the current time-step, while the initial value for nodes that do not exist in the previous step can be obtained by spatial interpolation on the parent grid. Concerning the second issue, two different cases are possible. As the first case, it is possible that a boundary of the refined mesh coincides with a physical boundary. In this case, the boundary condition for physical boundary was applied. However, if a boundary was located in the interior of the physical domain, the Dirichlet boundary condition was imposed using the interpolant produced by temporal interpolation between the initial and end point in the current interval. The procedure of mesh refinement continued until satisfying accuracy obtained at all nodes.

#### 2.5.2 Temporal discretization and solving stiff ODEs

After semi-discretization of the PDE system in space, an ordinary system of differential equations (ODE) was obtained. The resulted ODE system was solved using two different methods with adaptive time-steps. The first method, a Rosenbrock-type method, treated the ODEs implicitly and belongs to a class of implicit solvers that replace non-linear system of equations by a series of linear systems and known as linearly implicit methods. The main scheme of an s-stage Rosenbrock method for a non-autonomous system of ODEs, *My*’ = *f*(*t*, *y*), is as follows:

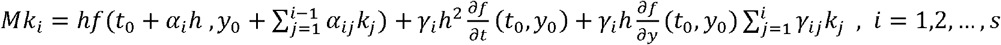

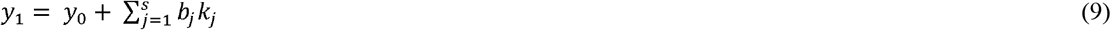

where *h* = *t*_1_ − *t*_0_ is the current time-step, b_j_, α_ij_ and γ_ij_ are coefficients chosen such that the order and stability conditions satisfied and 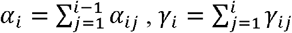.

The stiff ODEs were also solved by exponential integrators a class of numerical methods developed mainly to solve semilinear problems. Semilinear equations are a class of stiff differential equations with a stiff linear part and nonstiff nonlinear part. Reaction-diffusion equations constitute a prominent class of semilinear equations, where stiffness arises from linear term, namely diffusion, while the reaction term is usually nonlinear and non-stiff or mildly stiff. Discretization of the equation (1) leads to the following system of equations:

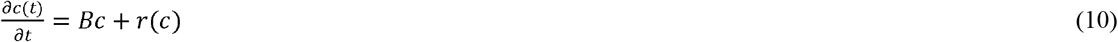

where *Bc* is a discretized approximation of *DΔc.* To obtain the solution at time t_n+1_=t_n_+Δt, integrating the above equation over time-step Δt after multiplication by the integrating factor *e^−Bt^* yields the following equation:

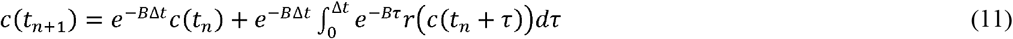

By solving the integral in the right hand side of the above equation using quadrature, c(t_n+1_) can be obtained. By this procedure the linear term is calculated exactly, while the nonlinear term is treated explicitly. Using different quadrature rules leads to different exponential integration schemes. In this work, exponential Runge-Kutta (EXPRK) scheme was used to solve the above equation (for more details about EXPRK implementation and exponential integration see [23]).

Selecting either LUMR or NUMR method for discretization and either exponential EXPRK or Rosenbrock method for ODE solving result in four different methods which compared in section 3.1: LUMR and EXPRK (LUMEX), LUMR and Rosenbrock method (LUMROS), NUMR and EXPRK (NUMEX), NUMR and Rosenbrock method (NUMROS).

#### 2.5.3 Hybrid stochastic-deterministic modeling

To model stochastic gating of IP_3_R channels, Gillespie method was used using the next reaction method for generating random numbers [24]. According to the method, it is assumed that there are m stochastic reactions (R1, R2, …, Rm) in the system, each associated with a propensity function (ai) determining the rate of that reaction. Per each reaction, a random variable r is drawn from a uniform distribution in the interval [0,1] and, then, the reaction with the smallest is selected to be executed. Subsequently, the number of molecules and propensity functions are updated considering Ri is executed. New random variables for the reactions with changed propensity functions are generated as mentioned and the process continues in the same manner. However, to couple the stochastic gating of IP_3_R channels with deterministic reaction-diffusion equations, some modifications are required, because the Ca2+ concentration is not constant during stochastic time steps due to the deterministic variations by reaction-diffusion events and/or channel gating. To solve this issue, hybrid stochastic-deterministic modeling approach was used as described in [25]. Briefly, per each reaction, a new variable gi (i=1,2,…,m) was considered with the following definition:

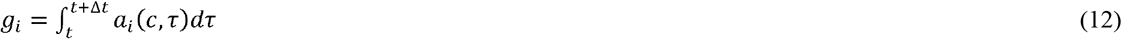

In addition, random variables ξ_*i*_ (*i*=*1*,*2*,…,m) were generated by drawing random numbers *r*_*i*_ from uniform distribution in the interval [0,1] and letting ξ_*i*_=*ln*(*1*/*r*_*i*_). The reaction *R*_*i*_ was executed whenever *g*_*i*_=ξ_*i*_ satisfied. From an implementation point of view, the following differential equations were considered and solved numerically along with the reaction-diffusion equations (deterministic part of the system):

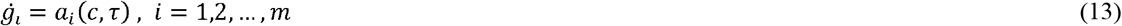

At the end of each time step, *g*_*i*_ values compared to their corresponding ξ_*i*_ random variables. Once *g*_*i*_=ξ_*i*_ satisfied for a reaction *R*_*i*_, the reaction executed, a new random variable ξ_*i*_ was generated and *g*_*i*_ was set equal to zero for that reaction.

All calculations and simulations were performed in MATLAB. Since each stochastic event was modeled by a random number, each simulation was repeated at least three times to avoid overestimating rare events.

## 3 Results and Discussion

### 3.1 Comparing numerical methods

In this section, discretization methods and ODE solvers described in sections 2.5.1 and 2.5.2 were compared in terms of their efficiency, namely the time of computation and memory usage, while maintaining their accuracy at the same level using temporal and spatial adaptivity.

Regarding the time of computation, two factors must be scrutinized: time consumption of computing each time-step and the length of time-steps. Due to spatial and temporal adaptivity, these two factors vary over simulation. To consider various events during the simulation, a sufficiently large time interval (10 ms) was simulated containing a single channel opening-closing event lasting for approximately 2 ms (figure 1). Based on the results, NUMEX algorithm took such a diminutive time-steps so that it was impossible to utilize this method to simulate a 10 ms time interval. On the other hand, LUMEX algorithm performed remarkably more efficient than both LUMROS and NUMROS, while comparing LUMROS and NUMROS algorithms denotes that the former was significantly faster.

Comparing the results for NUMEX and LUMEX methods, it can be inferred that the ODE system arising from NUMR discretization is considerably stiffer than the ODE system arising from LUMR discretization which stems mainly from the fact that non-uniform meshes often lead to stiffer problems.

**Figure 1:**
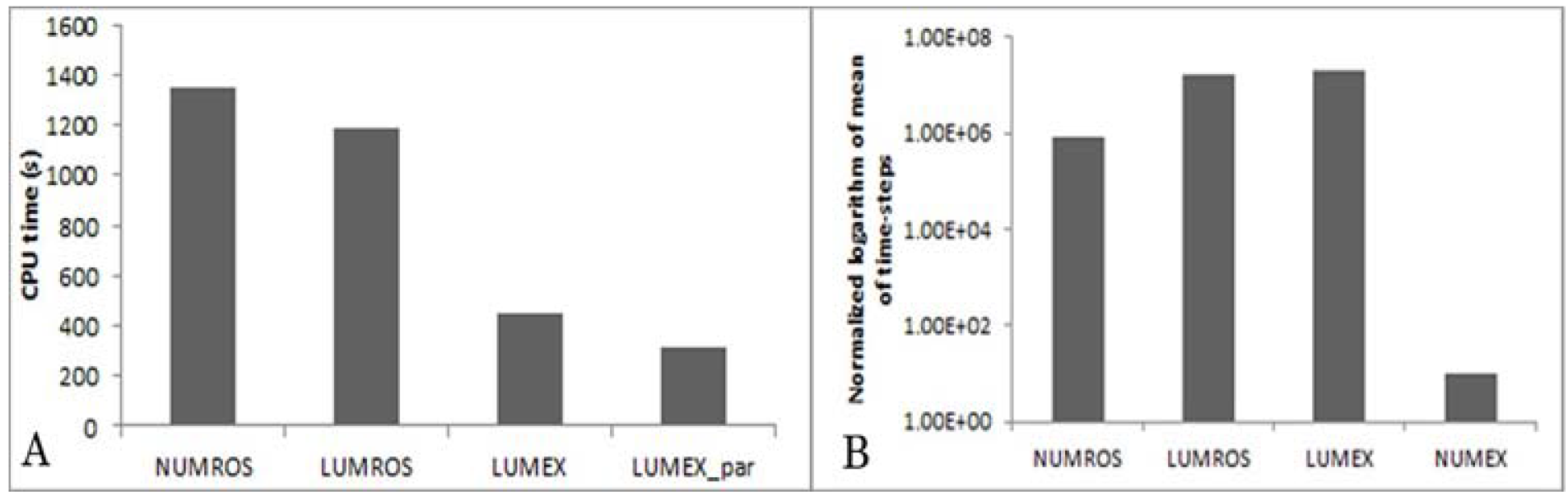
Comparing various numerical methods in terms of (A) CPU time and (B) mean time-step simulating 10 ms duration containing a single channel gating event.

Although fully implicit Rosenbrock method can be utilized in combination with both LUMR and NUMR gridding, its usage with LUMR enhanced the speed of computation with respect to the previously used NUMROS method. Since the subgrids with the same level of refinement are treated independently in the LUMR algorithm, parallelizing computation for this algorithm can be performed easily without introducing new error sources. By parallelizing LUMR, the time of computation even further decremented.

Comparing the two ODE solvers, Rosenbrock method has better stability properties and can be used for a more diverse range of problems, particularly stiff ODEs with stiff non-linear terms, while EXPRK, when applicable, can solve the problem more efficiently.

Although for this type of problem ameliorating the calculation speed is more important, the memory usage of the discretization algorithms was compared as memory might be the circumscribing factor when the size of the problem increases either through the size of the domain or the number of participating molecular species. Most of the memory usage of the algorithms is due to saving the solutions of the previous time-point on each grid, which is required to obtain the solution at the current time-point. This implies that the memory usage is mainly depends on the number of the nodes used to discretize the physical domain in each algorithm. Maintaining the same level of spatial error, the maximum number of required nodes to discretize the defined domain for LUMR was 20592 versus only 8050 for NUMR. Nevertheless, only a small portion of these nodes is required at each step of LUMR algorithm and memory usage can be optimized using memory management methods [22].

### 3.2 Electrophysiological modeling

To verify electrophysiological model of MSCs, the behavior of ionic currents passing through plasma membrane was modeled, while maintaining the membrane potential constant at steady-state (figure 2-A). The results are in accordance with patch clamp results obtained in the previous work [12] for the third group of MSCs denoted as “type C”. Furthermore, the effects of various types of ES applied to MSCs to stimulate differentiation on the membrane potential were investigated (figure 2-B).

**Figure 2:**
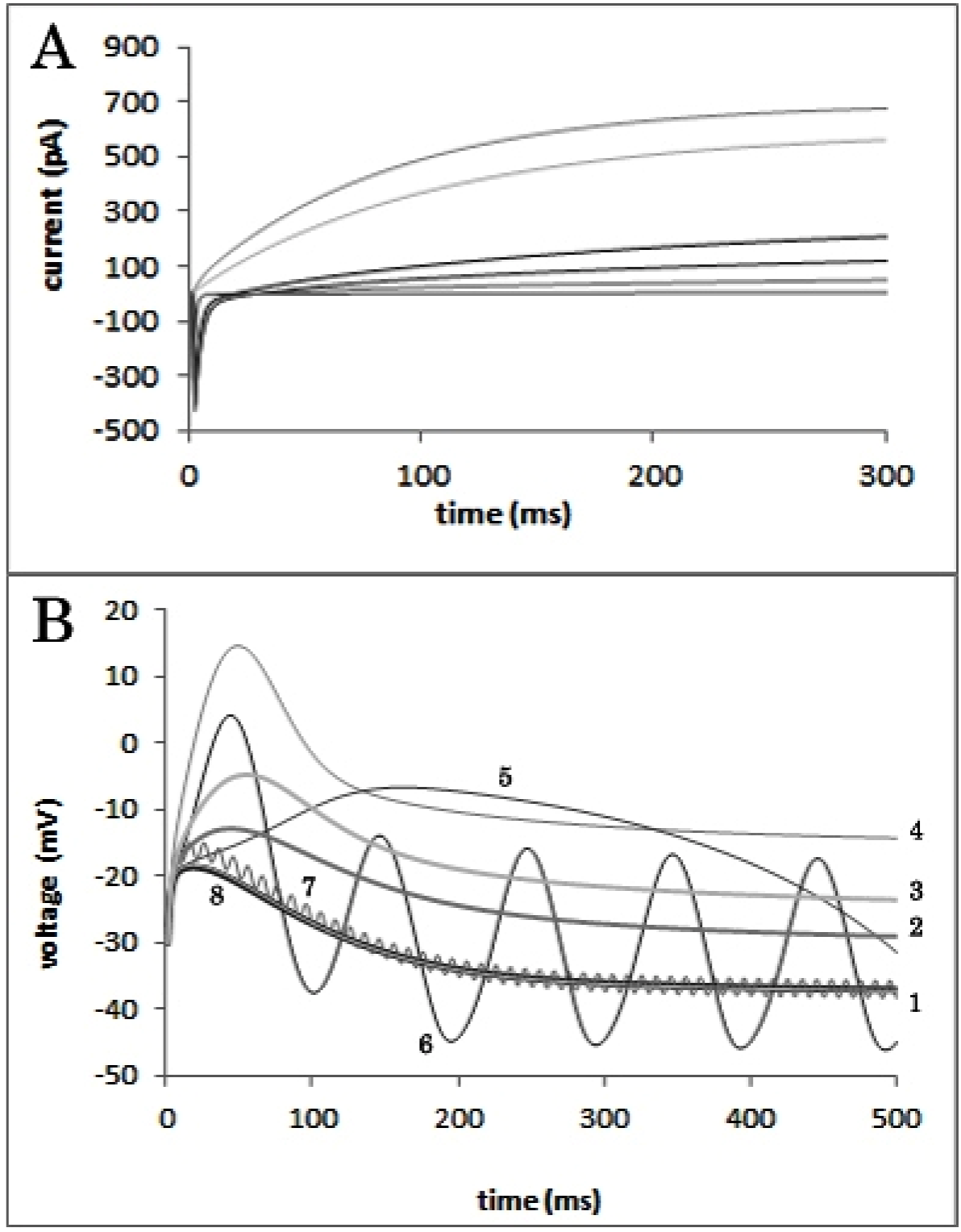
(A) Total ionic current passing through the plasma membrane of MSC type “C” at steady-state while maintaining the membrane potential constant. Negative and positive currents represent inward and outward direction, respectively. (B) The effect of various electrical stimulation regimens on the membrane potential. Each number in the diagram represents a special stimulation setup in the following manner: (1) no stimulation, DC stimulations with (2) 10 pA, (3) 20 pA, (4) 50 pA amplitudes and AC stimulations with 50 pA amplitude and (5) 1 Hz, (6) 10 Hz, (7) 100 Hz, (8) 1000 Hz frequencies.

Despite previously proposed hypothesis that ES triggers LCC opening by membrane depolarization, only DC and low frequency AC stimulations caused significant depolarization in the membrane potential. Taking into account that high frequency AC stimulations subtly alter the membrane potential might plausibly explicate the experimental observation that AC stimulations with less than 100 Hz frequency, particularly less than 30 Hz, are effective for tissue engineering purposes [26, 27]. The fact that there are several reports on the differentiating effect of pulsatile, biphasic and high frequency stimulations on MSCs which are mediated through LCCs [3, 28–30] implies that they influence LCC gating by another mechanism probably mediated through their frequency, as a shift in gating mode of LCC has been observed in response to the frequency of the stimulation [31]. Unfortunately, the kinetics of this modal shifting, particularly for higher frequencies, is poorly understood and more experimental investigations are needed to scrutinize the relationship between this modal shifting and ES effect on MSCs.

### 3.3 The stimulation effect on LCC and IP_3_R channels gating

In this section, the effects of ES on LCC and IP_3_Rs gating were scrutinized mediated respectively through membrane depolarization and Ca^2+^ diffusion into the cell supposing that Ca^2+^ entered to the subspace via single LCC.

The effect of various ES types with notable influence on the membrane potential (cf. Section 3.2) on LCC gating was evaluated in terms of the number of transitions to the long-lasting opening mode (figure 3-A). Notwithstanding DC stimulation with lower amplitude (20 pA), applying high amplitude DC stimulation (50 pA) increased the number of transitions considerably which is mostly due to the fact that LCCs steeply inclined toward the long-lasting opening mode by depolarization beyond 0 mV [16].

**Figure 3:**
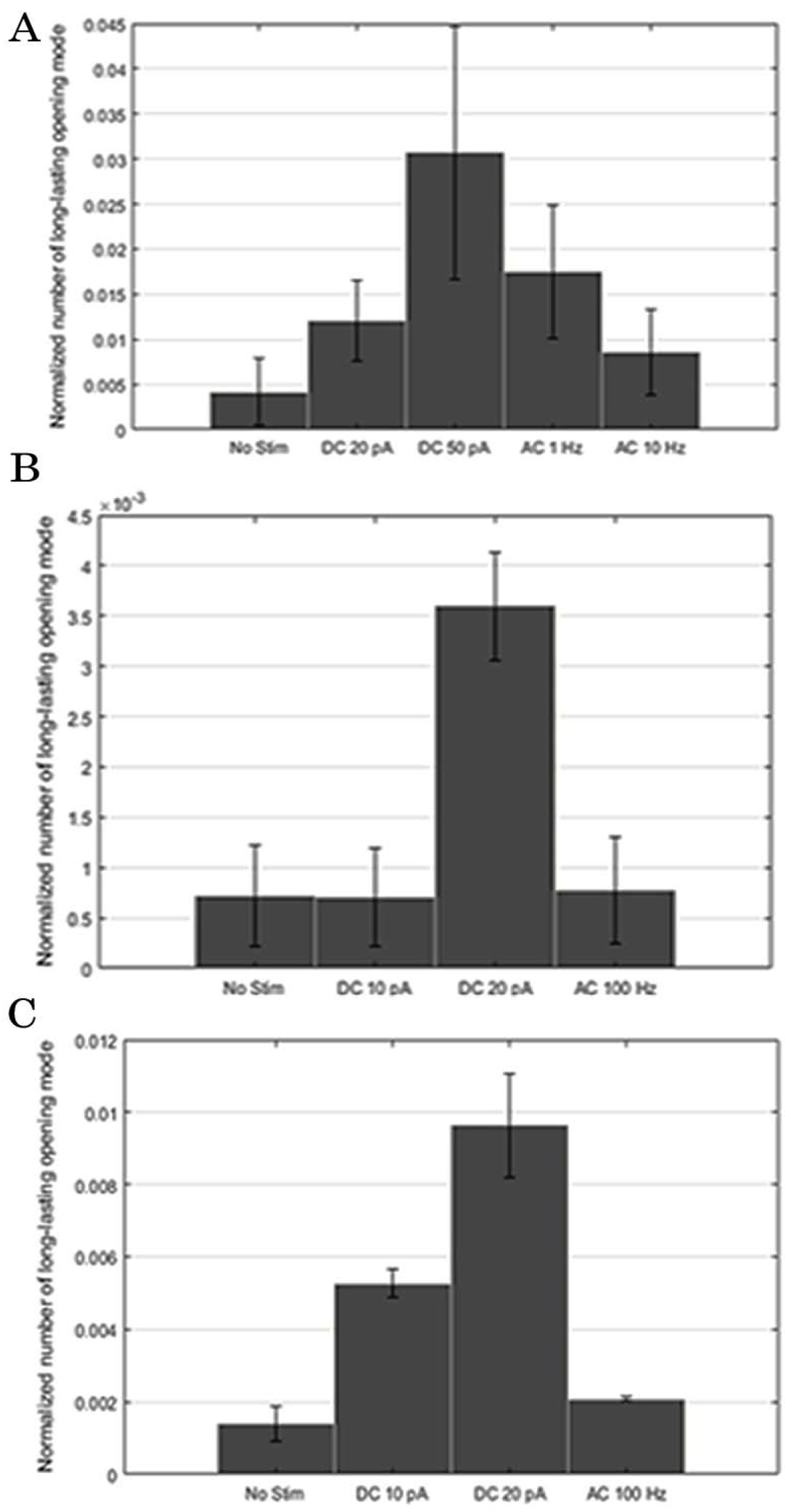
The number of transitions to the long-lasting opening mode in response to various electrical stimulation groups considering (A) single LCC, (B) four LCCs and (C) nine LCCs. In the diagram, “No Stim” represents no stimulation group and the amplitude of all AC stimulations is 50 pA. The results were represented as mean ± standard deviation.

Considering AC stimulation, the number of transitions to long-lasting mode depended on the frequency of stimulation and lower frequencies caused more transitions. On the other hand, high variation in the number of transitions was observed due to high variation in the membrane potential, especially AC stimulations with lower frequencies.

Evaluating ES impact on IP_3_R channels, the number of open channels in the cluster at each time during simulation was compared between no stimulation group and 50 pA DC stimulation and AC stimulation with 1 Hz frequency and 50 pA amplitude (figure 4). These two types of ES setup were selected as representatives of DC and low-frequency AC stimulation, respectively, due to their highly distinctive impact on the membrane potential and single LCC gating. The results indicated that applying ES (and hence more Ca^2+^ influx via LCC) caused the lower number of synchronous opening of IP_3_R channels and particularly, simultaneous opening of higher number of channels which can initiate local upsurge in Ca^2+^ concentration (Ca^2+^ puffs) significantly reduced with respect to no stimulation. This outcome is in accordance with a previous experimental study in which it was demonstrated that applying ES to MSCs reduced the number of local Ca^2+^ elevations [11].

**Figure 4:**
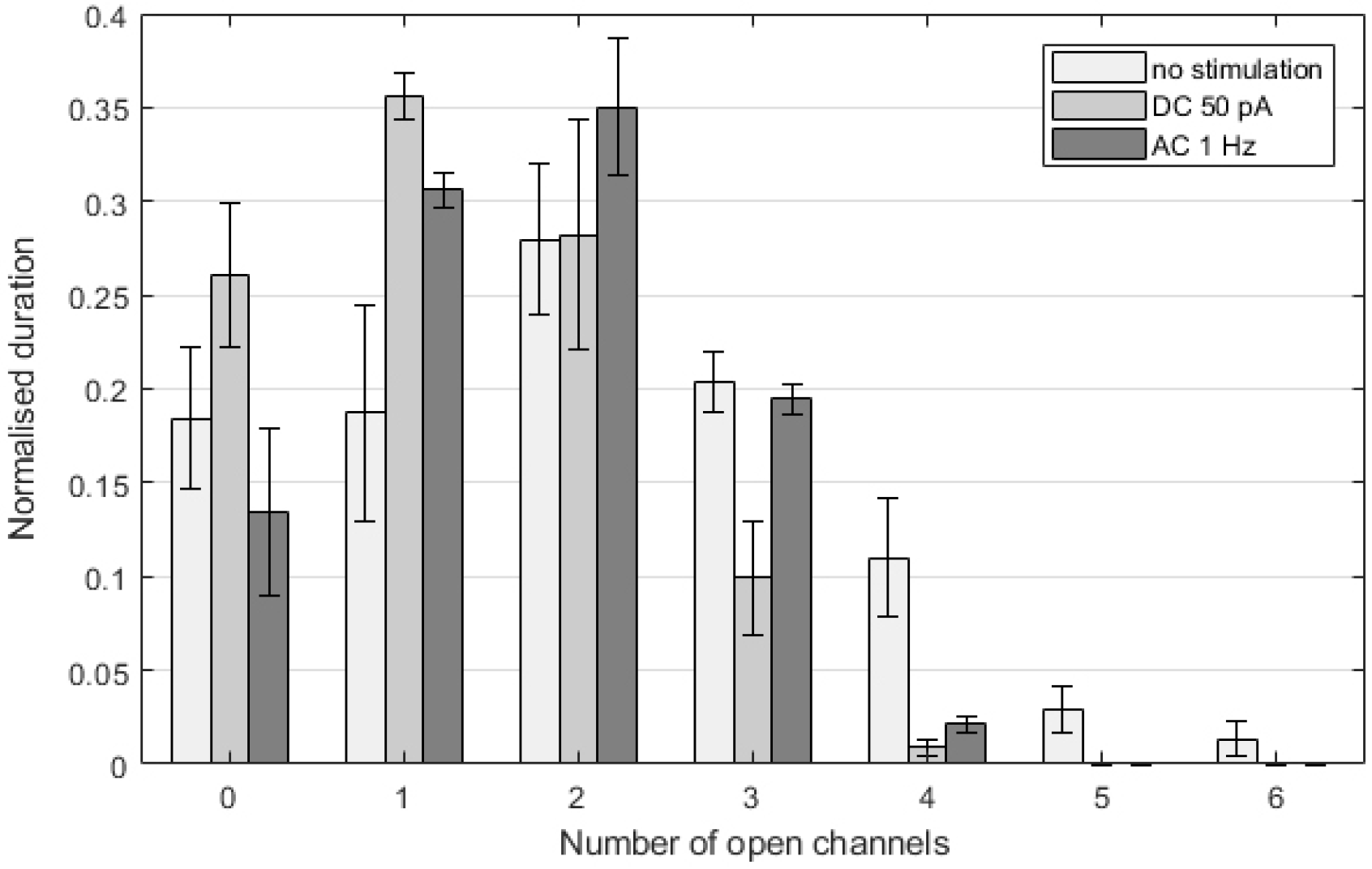
Normalized duration of various states of the cluster featured by the number of open channels. The electrical stimulation utilized for this simulation are 50 pA DC stimulation and AC stimulation with 50 pA amplitude and 1 Hz frequency.

### 3.4 The impact of multiple LCCs

Hitherto, all simulations were performed supposing that Ca^2+^entered into the subspace via only one LCC. As the open probability for single LCC rises considerably by depolarization beyond 0 mV [16], it is expected low amplitude DC stimulations, with subtle effects on the membrane potential, exert no effect on MSCs. However, the previous experimental results indicate even such stimulations reduce the number of local Ca^2+^ elevations in MSCs [11]. To provide a possible explication for this observation, it was assumed that Ca^2+^ can enter into the subspace via multiple LCCs and the effect of low amplitude DC stimulations on gating of multiple LCCs was scrutinized in this section (figure 3-B, C). Assuming Ca^2+^ entrance into the subspace through multiple LCCs, even small alteration in the membrane potential exerted distinguishable effect on Ca^2+^ influx which was more perceptible as the number of LCCs increased. Furthermore, the probability of transition to the long-lasting opening mode was less susceptible to stochastic variation supposing the higher number of LCCs. On the other hand, applying AC stimulation with lower frequency (100 Hz) had no special effect on modal-shifting even considering the higher number of LCCs which indicates that AC stimulations, especially with high frequencies, exert their effect on the cells via another mechanism as discussed in section 3.2.

It is rational to suppose that LCCs are distributed in plasma membrane such that Ca^2+^ entering via multiple LCCs can influence Ca^2+^ release from the ER and hence, efficaciously affect intracellular Ca^2+^ dynamics which is a possible explanation for the effect of low amplitude DC stimulations, although more evidence is required.

## 4 Conclusion

With growing interest in spatio-temporal modeling of intracellular events, especially signaling events in order to decode the spatial information of the signals, developing efficient numerical methods has become essential. In this article, a novel numerical method belonging to locally uniform mesh refinement, a class of numerical solvers, was proposed and its better performance with respect to several alternative methods was demonstrated. According to the benchmark comparison performed in this work, the incipiently proposed method was efficient for simulating intracellular Ca^2+^ spikes in the order of a few seconds comparing with a few milliseconds for previous methods [20, 32].

The model was used to investigate the effect of various ES setups which are used to differentiate stem cells for tissue engineering, particularly focusing on DC and AC stimulations with different amplitudes and frequencies. The results indicated that only certain DC stimulations with amplitudes higher than a threshold and AC stimulations with frequencies lower than a threshold can significantly alter the membrane potential and hence, trigger voltage-dependent LCC modal shifting. Although the effect of low-amplitude DC stimulations on the Ca^2+^ influx can be discerned from no stimulation assuming Ca^2+^ entrance via multiple LCCs, no change was observed by incrementing the number of LCCs for high frequency AC stimulations implying they exert their effect on the cells through another mechanism.

The model betokened that more transitions of LCCs to long-lasting opening mode, reduced the number of synchronous opening of IP3R channels in accordance with previous experimental results [11]. Besides, it was demonstrated the number and distribution of LCCs play a major role in determining the cellular response to ES. In general, it is inferred from the results that exploiting methods to increase LCC expression before applying ES is propitious as both the sensitivity of the cells increases and stochastic variation between different cells decreases. Both of these features might be consequential in tissue engineering, especially when ES is used to drive stem cells into different cell lineages in a cell culture, an essential requisite to construct organs in vitro situation.

More importantly, these findings bolster the applicability of such computational models in tissue engineering to enhance the current methods and fine-tune the stimuli to control cellular fate.

